# Functional connectivity alterations of the temporal lobe and hippocampus in semantic dementia and Alzheimer’s disease

**DOI:** 10.1101/322131

**Authors:** Simon Schwab, Soroosh Afyouni, Yan Chen, Zaizhu Han, Qihao Guo, Thomas Dierks, Lars-Olof Wahlund, Matthias Grieder

## Abstract

The severe semantic memory impairments in semantic dementia have been attributed to a pronounced atrophy and functional disruption of the anterior temporal lobes. In contrast, the medial and posterior temporal lobe damage predominantly found in patients with Alzheimer’s disease has been associated with episodic memory disturbance. However, the two dementia subtypes share hippocampal deterioration, despite a relatively spared episodic memory in semantic dementia. To gain more insight into the mutual and divergent functional alterations seen in Alzheimer’s disease and semantic dementia, we assessed the differences in intrinsic functional connectivity between temporal lobe regions in patients with Alzheimer’s disease (n = 16), semantic dementia patients from two international sites (n = 23), and healthy controls (n = 17). In an exploratory study, we used a functional parcellation of the temporal cortex to extract time series. The Alzheimer’s disease group showed a single connection with reduced functional connectivity as compared to the controls. This connection was located between the right orbitofrontal cortex and the right anterior temporal lobe. In contrast, functional connectivity was decreased in the semantic dementia group in six connections, mainly involving the hippocampus, lingual gyrus, temporal pole, and orbitofrontal cortex. We identified a common pathway with semantic dementia, since the functional connectivity between the right anterior temporal lobe and the right orbitofrontal cortex was reduced in both types of dementia. This might be related to social knowledge deficits as part of semantic memory decline. However, such interpretations are preferably made in the context of all disease-specific semantic impairments and functional connectivity changes. Despite some limitations owed to the two database sites, this study provides a first preliminary picture of the brain’s functional dysconnectivity in Alzheimer’s disease and semantic dementia. Future studies are needed to replicate findings of such a common pathway with matched diagnosis, neuropsychological, and data MRI acquisition procedures.

## Introduction

Everybody occasionally experiences difficulties in integrating past events into an accurate context — a condition classified as an episodic memory disturbance. Intact episodic memory [1] requires the processing of information about chronology, place and the protagonists who were involved in an event. The capability to store and retrieve autobiographical memory is however not sufficient for an intact episodic memory. Humans also strongly rely on a fully functioning semantic memory. Concretely, semantic memory reflects our general knowledge about concepts such as objects, people, and words. Thus, only a sound interplay of these two memory systems, episodic and semantic memory, allows a cognitively healthy state of an individual.

Previously two initially contradicting models of the neurophysiological organization of semantic memory have been harmonized as what can be characterized as a ‘cortically distributed plus semantic hub’ theory [2,3]. The term “distributed” refers to the idea that regions which process semantic concepts receive multimodal input from corresponding brain regions (e.g., visual attributes from visual brain regions, tactile attributes from the sensorimotor cortex, etc.). Subsequently, these multimodal inputs from distributed cortical areas converge to so-called unitary semantic concepts in the semantic hub [4,5]. The semantic hub was found to be localized bilaterally in the anterior temporal lobe, a region which is atrophied and hypometabolized in patients with the semantic variant of primary progressive aphasia, also known as the temporal variant of frontotemporal dementia (FTD) or semantic dementia (SD) [5–7]. In SD, the onset of gray matter atrophy occurs in the anterior temporal lobes, frequently with an asymmetry towards the more affected left hemisphere. With progression of the disease, the temporal pole and medial as well as lateral temporal areas are degenerated [8]. However, the patients seem to exhibit an almost intact episodic memory, when tested non-verbally, while their semantic memory is severely deteriorated [9,10].

In contrast to SD, patients with Alzheimer’s disease (AD) show predominantly episodic memory impairments, and semantic memory deficits can only be observed to a minor degree [11–13]. AD has been described as a disconnection syndrome, that is, connections of functionally or structurally linked brain regions that are part of a network become increasingly disrupted [14–16]. This degenerative mechanism has been associated with the cognitive deficits of patients with AD [17–19]. A common finding in AD is that gray matter atrophy onset can be localized in the hippocampal, posterior cingulate and lateral parietal brain regions, as well as in the amygdala [20,21]. The hippocampus forms a core region for episodic memory encoding. However, it has also been associated with semantic memory functions [22]. In fact, Burianova and colleagues [22] postulated that the hippocampus is part of a *common* declarative memory network, suggesting that the hippocampus has a key role in both semantic as well as episodic and autobiographical memory.

The properties of functional systems, as for example Burianova and colleagues’ proposed declarative memory network, are commonly assessed by the use of a resting-state functional connectivity (FC) analysis. The human resting-state is characterized by spatially discriminate brain regions that co-activate and deactivate at a low temporal frequency, commonly known as resting-state networks [23,24]. These functional systems, or resting-state networks, are commonly assessed using blood-oxygen level dependent resting-state fMRI. It has become very popular to study FC alterations in various mental and neurological disorders including AD, demonstrating a relationship between disease and abnormalities in resting-state networks [25–27].

FC changes (i.e., decreases and increases of connectivity strengths) in AD have been found predominantly in the hippocampus and the default mode network [28–31]. With the progression of the disease, structural and functional connectivity distortions affect several networks, particularly those involving the parahippocampus [17,32]. In SD, FC appears to be deteriorated in regions either affected by or proximate to the core of atrophy, located in regions such as the temporal pole, anterior middle temporal gyrus, inferior temporal gyrus, and insula [6,33–35]. Furthermore, reduced FC of the anterior temporal lobe with various cortical regions was also found in SD [2].

Considering these findings as well as the distinct pathology of AD and SD, it is likely that the neuronal loss of hippocampal cells that results in gray matter atrophy certainly affects the functional networks in a way that generates episodic memory deficits. Temporal pole atrophy alone might not be necessary (but sufficient) to lead to semantic impairment. Following these findings, La Joie et al. [36] identified the hippocampus as the ‘main crossroad’ between brain networks that are disrupted in AD and SD. Despite the growing body of research, the common and divergent changes of FC among regions of the temporal lobes in AD and SD are not fully understood. A caveat when interpreting the existing literature is the common use of anatomical/structural parcellation instead of a functional parcellation to study FC. Functional parcellations have the advantage that the resulting functional ROIs are homogeneous, i.e., the voxels have similar time courses. On the other hand, parcellations based on brain structure can merge the time series across functionally different areas which can be problematic [37].

This proof-of-concept study aimed at disentangling FC alterations of the temporal lobe in AD and SD using a refined division of temporal subregions: sixty-six functional regions of interest (ROIs) of the temporal lobes from a functional atlas [38]. In contrast to numerous previous studies, we accounted for structural changes (i.e., gray matter atrophy) in order to extract FC time series data from preserved gray matter tissue which can still be functional [39,40]. In other words, results from the FC analysis reflect the functional reorganization of the temporal lobes affected by atrophy.

A common issue with studies involving patients with SD is the small sample size due to the low prevalence and relatively difficult diagnosis. In order to overcome this to some extent, we pooled two data sets from two different recording sites (see Method section for details). Orban et al. [41] showed in multivariate fMRI analysis the advantage of multisite fMRI-data. Their approach appears to be generalizable; however, we will address some of its limitations in the discussion section (e.g. site-specific diagnostic criteria, neuropsychological testing, and fMRI acquisition procedures).

Despite the exploratory character of the study, based on previous findings described above, the following hypotheses were tested: in AD, we expected FC alterations in the hippocampus, parahippocampal ROIs, and possibly posterior temporal ROIs. In SD, altered FC was anticipated in the hippocampus, the fusiform gyrus, and the temporal pole.

## Methods

### Participants

We analyzed resting-state fMRI data from a total of 62 participants from three groups: semantic dementia (SD), Alzheimer’s disease (AD), and a healthy elderly control group (HC). We examined all the functional MRI data and excluded six datasets due to insufficient data quality (see data quality control). The final sample consisted of 56 participants: Twenty-three patients with SD, with a mean age (± standard deviation) of 62 ± 7.6, 16 patients with AD, mean age of 70 ± 8.5, and 17 individuals in the HC group, mean age 70 ± 3.4; see Table 1 for demographics and clinical variables. Patients with SD from the Stockholm site (n = 7) were recruited throughout Sweden and diagnosed using the criteria of Neary et al. [42], while patients with SD from Shanghai were recruited from Huashan Hospital in Shanghai (n = 19), according to the criteria of Gorno-Tempini et al. [43]. The main diagnostic criteria of both guidelines share clinical observation features such as impaired word naming and comprehension, spared repetition, and surface dyslexia and dysgraphia. Differences in these two diagnostic criteria, as for instance the introduction of brain imaging as a supportive diagnostic feature in Gorno-Tempini et al. (2011), were not relevant, because also the Swedish patients underwent MRI to assess anterior temporal lobe atrophy. Patients with AD were recruited at the Memory Clinic of the Geriatric Department at Karolinska University Hospital in Huddinge, Sweden (n = 19). Their diagnosis was performed by expert clinicians and was in accordance with the ICD-10 criteria [44]. The patients with AD included in this study underwent a standard clinical procedure which consisted of examinations such as structural neuroimaging, lumbar puncture, blood analyses, and a neuropsychological assessment (these assessments were part of the clinical routine and only used for diagnosis). Further inclusion criteria for patients from the Stockholm site was a Global Deterioration Scale lower than 6 (i.e., moderate dementia or milder) and the Cornell Depression Scale below 8. Healthy elderly controls were recruited by advertisement (n = 22) in the Stockholm area. Presence of medical or psychiatric disorders (other than dementia), intake of drugs affecting the nervous system, or magnetic implants, led to an exclusion from the study. Variables available for all participants included in the study were age, gender, and Mini-Mental State Examination (MMSE).

**Table 1.** Descriptives and clinical scores. Kruskal-Wallis tests were run to assess group differences of age, education, MMSE, BNT, lexical decision, AF, and VF. Comparisons between AD and SD of the CDS and GDS scores were performed using the non-parametric Kolmogorov-Smirnov-Test.

|  | HC (n=17) | Normative data<sup>†</sup> | AD (n=16) | SD (n=23) |  |
| --- | --- | --- | --- | --- | --- |
|  | Mean (std. dev.) | Mean (std. dev.) | Mean (std. dev.) | Mean (std. dev.) | p-value |
| Age, years | 67.9 (3.3) |  | 68.4 (8.5) | **61.5 (7.4)** | 0.004 |
| Gender (F:M) | 12:5 |  | 7:9 | **10:13** | - |
| Education, years | 13.9 (3.1) |  | 13.1 (3.0) | 12.4 (1.5)<sup>3</sup> | 0.61 |
| CDS | - |  | 1.0 (1.0) | 1.8 (2.2)<sup>3</sup> | 0.60 |
| GDS | - |  | 2.9 (0.8) | 3.8 (0.4)<sup>3</sup> | 0.031 |
| MMSE (max 30) | 28.8 (0.8) |  | 24.5 (4.8) | **20.8 (5.2)<sup>4</sup>** | < 0.0001 |
| BNT (max 60) | 54.4 (3.7) | 54.0 (4.5) | 45.6 (6.5) | 8.2 (5.7)<sup>3</sup> | < 0.0001 |
| Oral picture-naming (max 140) | - |  | - | 39.2 (27.6)<sup>5</sup> | - |
| Word-triple association (max 70) | - |  | - | 51.2 (10.1)<sup>5</sup> | - |
| Lexical decision (max 352) | 346.0 (3.7)<sup>1</sup> |  | 333.2 (23.5)<sup>2</sup> | 325.3 (23.0)<sup>6</sup> | 0.002 |
| AF, animals/minute | 23.8 (5.9) | 18.2 (3.8) | 14.1 (4.2) | 5.6 (4.3)<sup>3</sup> | < 0.0001 |
| VF, verbs/minute | 21.9 (5.8) | 18.2 (5.6) | 11.9 (5.0) | 7.0 (2.8)<sup>3</sup> | < 0.0001 |

<sup>†</sup> Normative data are reference values for comparison of the control group (HC) with respect to BNT with N = 32 [79]; AF with N = 94 [80]; VF with N = 67 [81]
<sup>1</sup> n = 16, <sup>2</sup> n = 12, <sup>3</sup> n = 5, <sup>4</sup> n = 19, <sup>5</sup> n = 14, <sup>6</sup> n = 4
Abbreviations: CDS = Cornell Depression Scale; GDS = Global Deterioration Scale; MMSE = Mini-Mental State Examination; BNT = Boston Naming Test; AF = animal fluency; VF = verb fluency.

All study participants provided informed consent prior to the data acquisition. The Shanghai study was approved by the Institutional Review Board of the State Key Laboratory of Cognitive Neuroscience and Learning, Beijing Normal University [33]. The Stockholm study was approved by the Regional Ethics Committee of Stockholm, Sweden.

### MRI data

MR images were acquired on two sites: The Karolinska Institute in Stockholm, Sweden, and the Huashan Hospital in Shanghai, China.

#### Stockholm site

MR images were acquired with a 3T Siemens Magnetom Trio scanner (Siemens AG, Erlangen, Germany). Structural images were 3D T1-weighted magnetization-prepared rapid gradient echo (MPRAGE) images using the following parameters: TR = 1900 ms, TE = 2.57 ms, flip angle = 9°, matrix size = 256 × 256, field of view = 230 × 230 mm2, slice number = 176 slices, slice thickness = 1 mm, and voxel size = 0.90 × 0.90 × 1 mm3. The structural images were previously used for voxel-based morphometry and published with a different purpose and sample configuration [45,46]. Functional images were acquired with a 32-channel head coil, using an interleaved EPI sequence (400 volumes; 26 slices; voxel, 3 ⨉ 3 ⨉ 4 mm3; gap thickness, 0.2 mm; matrix size, 80 ⨉ 80; FOV, 240 ⨉ 240 mm2; TR, 1600 ms; TE, 35 ms).

#### Shanghai site

Images were acquired with a 3T Siemens Magnetom Verio. Structural images were 3D T1-weighted magnetization-prepared rapid gradient echo (MPRAGE) images using the following parameters: TR = 2300 ms, TE = 2.98 ms, flip angle = 9°, matrix size = 240 ⨉ 256, field of view = 240 ⨉ 256 mm2, slice number = 192 slices, slice thickness = 1 mm, and voxel size = 1 ⨉ 1 ⨉ 1 mm3. Functional images were acquired with a 32-channel head coil, using an interleaved EPI sequence (200 volumes; 33 slices; voxel, 4 ⨉ 4 ⨉ 4 mm3; gap thickness, 0 mm; matrix size, 64 ⨉ 64; FOV, 256 ⨉ 256 mm2; TR, 2000 ms; TE, 35 ms, flip angle 90°). The data were previously published with a different sample configuration (SD only sample) in a combined structural and functional study using a hippocampus seed region [47], as well as in a structural VBM study [33].

### Preprocessing of functional MRI scans

We performed pre-processing using SPM12 (http://www.fil.ion.ucl.ac.uk/spm). We initially set all images’ origin to the anterior commissure, and then performed slice-time correction, realignment, coregistration, normalization to MNI space (2 ⨉ 2 ⨉ 2 mm3), and smoothing (full width half maximum [FWHM]; 8 mm). Time series data were high-pass filtered (1/128 Hz) and we regressed out 14 nuisance parameters (6 movement parameters and their first derivative, white matter, and CSF).

We carefully assessed data quality and inspected the spatio-temporal quality of each scan by comparing the slow and fast components of the data using DSE (Dvar, Svar & Evar) decomposition [48]. The DSE technique decomposes the dataset into three main components: fast, which is the squared mean difference; slow, which is the squared mean averages, and Evar, which refers to the sum of squares of the two ends of the time series. Subjects with remarkably high divergence (>75%-tile) between Dvar and Svar components were removed, as suggested in Afyouni & Nichols [48]. Therefore, we removed one SD and three HC datasets from the analysis. We further excluded two AD subjects, as more than 20% of their DVARS data-point were found to be corrupted. The remaining subjects were scrubbed as suggested by Power et al. [49]. Altogether, we excluded six datasets (9.7%) due to poor data quality. We re-run the diagnostics on the final sample and found no difference between groups regarding the DSE diagnostics (one-way ANOVA, all p > 0.05).

### Functional connectivity analysis

We investigated FC between each of the ROIs in three participant groups (AD, SD, and healthy elderly controls). We focused our analysis on the temporal lobes with the following rationale: first, brain regions identified as the origin of atrophy are located in the temporal lobe. Second, a ‘crossroad’ in FC network disruption in AD and SD was found in the hippocampus. Third, functional hubs for episodic and semantic memory can be found in the temporal lobe (as outlined above). Fourth, the strongest FC of temporal regions is located within the temporal lobes and concurs with functional networks crucial for language processing, the core clinical feature of SD [50]. The functional parcellation we used is based on resting-state fMRI data which was clustered into spatially coherent regions of homogeneous FC and was evaluated in terms of the generalizability of group level results to the individual [38]. From the 200 ROIs, we used a subset of 66 temporal ROIs that covered at least 5% or more of one of the following temporal structures from WFU Pickatlas 3.0.4 [51]: the superior temporal cortex, the middle temporal cortex, the inferior temporal cortex, the temporal pole, the hippocampus, the parahippocampal cortex, the lingual gyrus, the amygdala, the insular cortex, and the fusiform gyrus; these 66 ROIs are shown in Supplementary Figure 1. Analyzing merely 66 temporal ROIs leads to 2,145 pairwise correlations, which necessitates a strong adjustment for multiple comparisons to control for false positives. Using an even higher number of ROIs, for example comparing 200 ROIs in the whole brain, would require an even stronger correction (correcting for almost 20,000 comparisons). Such corrections would result in a sensitivity too low to detect even substantial FC changes.

We extracted the mean time series from the gray matter (probability > 0.70) of these ROIs to assure that time series are not contaminated with CSF signals from atrophied areas, resulting in 66 time series per subject. To address motion and physiological confounds which are global in nature, we applied global signal regression to the time series [52–54]. We created a pair-wise correlation matrix and transformed the correlation coefficient to Z-scores by Fisher’s transformation. We conducted a one-way ANOVA for each ROI pair (2,145 tests) to test the null hypothesis of no difference between the three groups. We also performed an additional analysis with age, mean gray matter probability in the temporal cortex and study site as additional covariates. Covariates can be problematic if these differ between groups [55], therefore, we performed sensitivity analyses with and without covariates and compared results. The ANCOVA with covariates produced the same results, i.e. the same connections had a statistically significant group effect. From the 2,145 total connections, we found 321 (sensitivity analysis: 324) significant edges that showed a group effect (uncorrected, *p* < 0.05), and after correcting the *p*-values for multiple comparisons, seven edges showed a significant group effect (FDR corrected, *p* < 0.05) in both the analyses.

### Voxel-based morphometry analysis

We additionally performed a voxel-based morphometry (VBM) analysis to quantify gray matter loss in the patients from the anatomical T1 images. VBM is a voxel-wise comparison of the local amount of gray matter volume between two groups [56]. We performed the following processing steps: spatial registration to MNI space (voxel size: 1.5 ⨉ 1.5 ⨉ 1.5 mm3) and tissue segmentation, bias correction of the intensities, smoothing of the GM images with 8 mm FWHM, and modulation by scaling with the total volume so that the resulting amount of gray matter in the modulated images remains the same as in the native images. In other words, this step removes the introduced bias from the registration of different brain sizes to MNI space. The Stockholm sample was registered using the European brain template, the Shanghai sample with the East Asian brain template and normalized to MNI space. We used the “Computational Anatomy Toolbox” CAT12 [57] and SPM12 [58] for the VBM analysis. Statistical inference was performed with the “Statistical NonParametric Mapping” software SnPM13 using non-parametric permutation/randomization two-sample *t*-tests with a voxel-wise family-wise error correction (FWE) of 0.05. We performed two *t*-tests and compared the HC group vs. the SD group, and the HC group vs. the AD group. Unlike in the analyses of the functional data where we excluded six datasets, the structural T1 scan from all the subjects were used in this analysis, the group sizes were HC with n = 20, AD with n = 18, SD with n = 24.

## Results

We first describe the clinical presentation of the patients included in this study (see Table 1 for details). The SD group performed poorer in MMSE than the AD group (Kruskal-Wallis over all groups: H = 29.5, df = 2, *p* < 0.0001; Kolmogorov-Smirnov group-wise post-hoc tests: HC-AD Z = 1.86, *p* = 0.002, HC-SD Z = 2.52, *p* < 0.001, AD-SD Z = 1.44, *p* = 0.033). Furthermore, the SD group showed significantly lower scores in the Boston Naming Test (BNT) than the AD group (Kruskal-Wallis over all groups: H = 23.3, df = 2, *p* < 0.0001; Kolmogorov-Smirnov group-wise post-hoc tests: HC-AD Z = 1.85, *p* = 0.002, HC-SD Z = 1.97, *p* = 0.001, AD-SD Z = 1.95, *p* = 0.001). Within the SD group, we observed that the impaired performance in picture naming (BNT, Stockholm site; oral picture-naming, Shanghai site) were more pronounced than lexical decision (Stockholm site) and word-triple association (Shanghai site), see Table 1. The group differences between SD and AD in MMSE and BNT are common findings given that the BNT is a semantic task and the MMSE relies on language comprehension, as both semantics and language are typically more affected in SD than AD. Finally, our AD group also showed semantic deficits as compared to the healthy control group (based on BNT, AF, and VF). MMSE was the only available neuropsychological test score for all participants from both sites, whereas the remaining tests were site-specific and therefore not comparable.

Next, we report the gray matter atrophy found in the patient groups, see Figure 1. In the SD patients (Figure 1A) we found two clusters of atrophy. The first was located in the left anterior medial temporal cortex, with a peak effect in the left temporal fusiform cortex (peak *t*-score = 14.0, *p*_FDR_ = 0.0021, df = 42; location at x = −34, y = −3, z = −36; cluster area 80.7 cm^3^). The second cluster was located in the temporal fusiform cortex of the right hemisphere (peak *t*-score = 10.6, *p*_FDR_ = 0.0021, df = 42; location at x = 34, y = −3, z = −34; cluster area 40.1 cm^3^). In the AD patients, we found two clusters with lower GM volume compared to controls in the left amygdala (peak *t*-score = 8.72, *p*_FDR_ = 0.006, df = 36; location at x = −26, y = −10, z = −12; cluster area 7.23 cm^3^) and the right amygdala (peak *t*-score = 7.49, *p*_FDR_ = 0.006, df = 36; location at x = 22, y = −3, z = −15; cluster area 7.47 cm^3^), see Figure 1B. A commonly expected hippocampal atrophy was yielded only with a more liberal threshold (Supplementary Figure 3).

**Figure 1:**
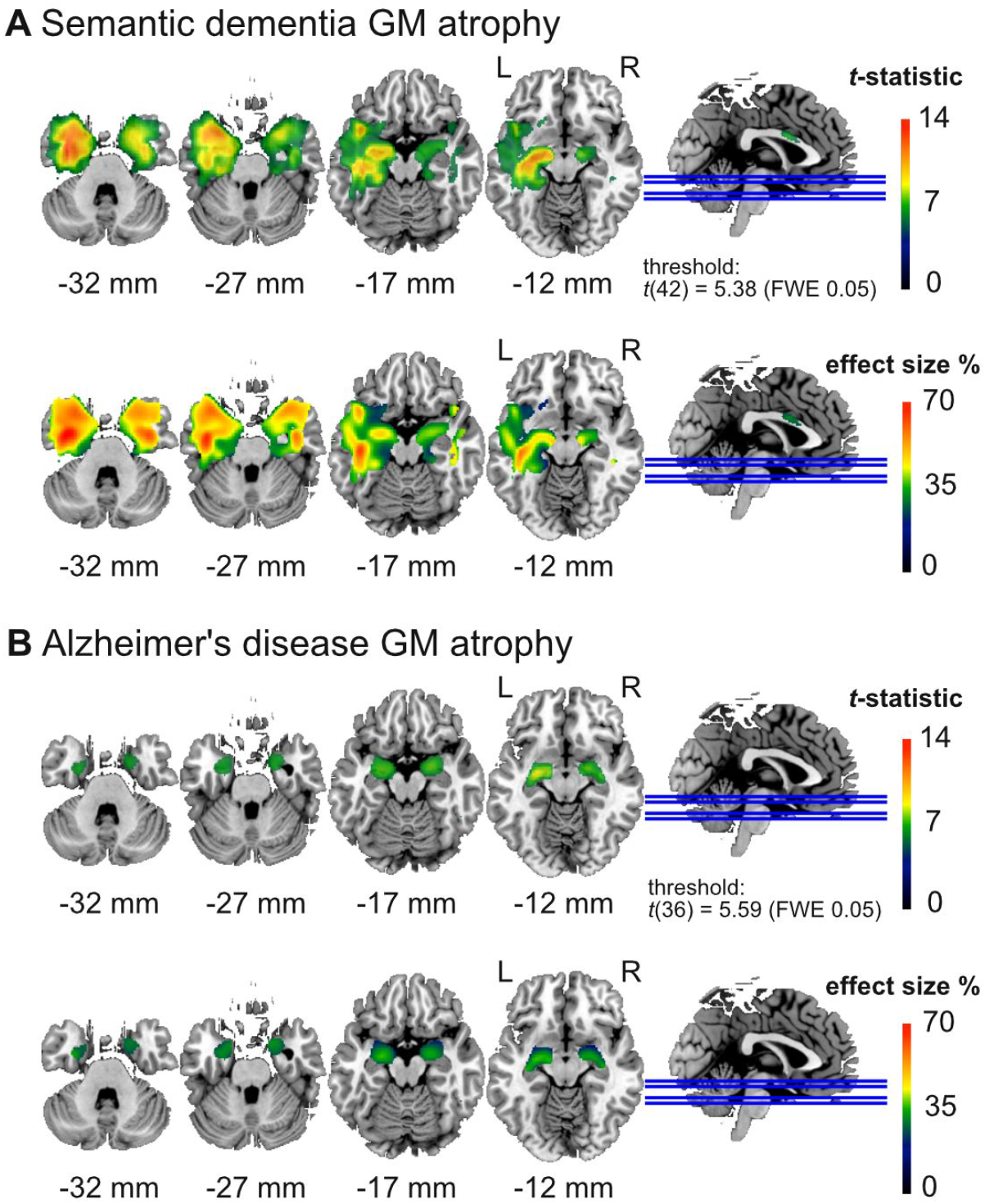
Areas with significant (voxel-level) gray matter (GM) atrophy (top) and effect size in terms of percentage GM reduction (bottom) in (A) the semantic dementia (SD) patients (n = 24) and (B) Alzheimer’s disease (AD) patients (n = 18) compared to the healthy elderly control group (n = 20). SD patients have atrophy in widespread areas of the left anterior temporal cortex including the temporal pole, while the AD patients show atrophy in the amygdala. SD patients showed more severe GM loss with up to 70% reduction, and AD patients with up to 40% reduction in some areas.

To achieve the main goal of this study, we analyzed the functional connectivity of 56 participants using 66 functional ROIs of the temporal cortex and related subcortical areas (see complete correlational matrix in Supplementary Figure 2). Seven connections (FC between ROI pairs) demonstrated a significant difference between the three groups after correcting for multiple comparisons (FDR corrected, *p* < 0.05). A detailed characterization and test statistics of these seven connections are shown in Table 2; the Z-values for the significant connections are depicted in Figure 2. We performed post-hoc tests (Tukey HSD) for single comparisons of the three groups to investigate the particular group contrasts that drove the significant group effect (Table 3). We found that most differences were related to the SD patients with significant changes in all of the seven connections. SD patients showed lower FC in 6 out of 7 connections compared to HC, and higher FC in one connection (edge no. 2) compared to HC. This higher FC in the SD patients was also significantly higher compared to the AD patients. The AD patients had a lower FC compared to HC patients in only 1 out of the 7 connections (edge 6). Comparing the two patient groups SD, versus AD, we found that SD had a significant lower FC in 4 connections (edges 3, 4, 5, 7).

**Table 2.** Seven functional connections that demonstrated significant group differences.

| Edge no. | ROI no. | ROI no. | Region | Region | F-value | FDR adj. p-value |
| --- | --- | --- | --- | --- | --- | --- |
| 1 | 129 | 11 | Left anterior superior temporal gyrus / middle temporal gyrus / insular cortex | Left posterior middle temporal gyrus / superior temporal gyrus | 10.86 | 0.034 |
| 2 | 85 | 24 | Right lateral inferior occipital cortex / lateral superior occipital cortex | Left posterior superior temporal gyrus / central opercular cortex / parietal opercular cortex / planum temporale | 11.52 | 0.030 |
| 3 | 198 | 32 | Right fusiform cortex / parahippocampal gyrus | Right inferior temporal pole | 12.64 | 0.026 |
| 4 | 70 | 37 | Left lingual gyrus / intracalcarine cortex / precuneus cortex | Left posterior hippocampus / thalamus | 13.18 | 0.026 |
| 5 | 89 | 37 | Right lingual gyrus / intracalcarine cortex | Left posterior hippocampus / thalamus | 10.95 | 0.034 |
| 6 | 153 | 71 | Right anterior middle temporal gyrus / superior temporal gyrus | Right orbitofrontal cortex | 12.42 | 0.026 |
| 7 | 112 | 72 | Left orbitofrontal cortex / insular cortex | Left anterior inferior temporal gyrus / middle temporal gyrus | 11.06 | 0.026 |

**Table 3.** Post-hoc Tukey HSD *p*-values for the three single comparisons (rows) and for each of the seven ROI-pairs that had a significant group effect (columns). Significant values reflect that the group effect was driven by a specific group level contrast.

| Edge No. | 1 | 2 | 3 | 4 | 5 | 6 | 7 |
| --- | --- | --- | --- | --- | --- | --- | --- |
| AD vs HC | 0.098 | 0.082 | 0.98 | 0.54 | 0.22 | 0.0005 | 1.00 |
| SD vs HC | < 0.0001 | 0.044 | 0.0002 | < 0.0001 | < 0.0001 | < 0.0001 | 0.0004 |
| SD vs AD | 0.064 | <0.0001 | 0.0004 | 0.002 | 0.024 | 0.97 | 0.0004 |

**Figure 2:**
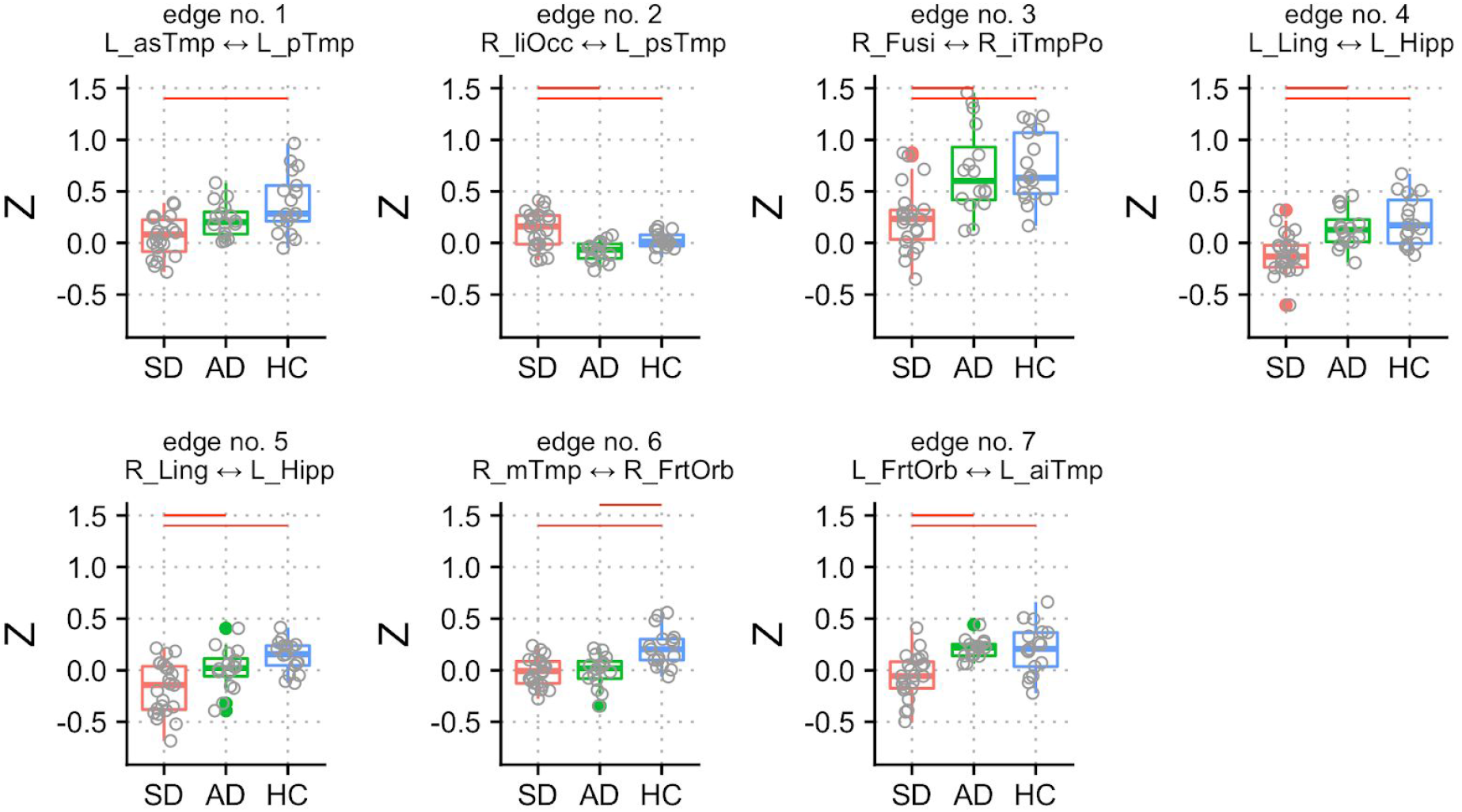
Z-scores of seven connections (edges 1–7) with significant group differences. Post-hoc tests between the three groups were performed, and significant group differences are denoted with red horizontal lines (see Table 2 for a detailed description of the ROIs). Gray circles represent single subject data points. Colored circles represent outliers.

We visualized the connectivity structure and connection strengths, see Figure 3. The SD patients generally had a much lower connectivity compared to the other two groups. An exception was the stronger contralateral connection between the right lateral inferior occipital cortex and the left posterior superior temporal gyrus (edge no. 2). The AD patients showed a lower FC compared to HC between the right middle temporal gyrus and the right frontal orbital cortex (edge no. 6). A common finding in all the three groups was that the FC between the right fusiform cortex and the right inferior temporal pole was the strongest (no. 3).

**Figure 3:**
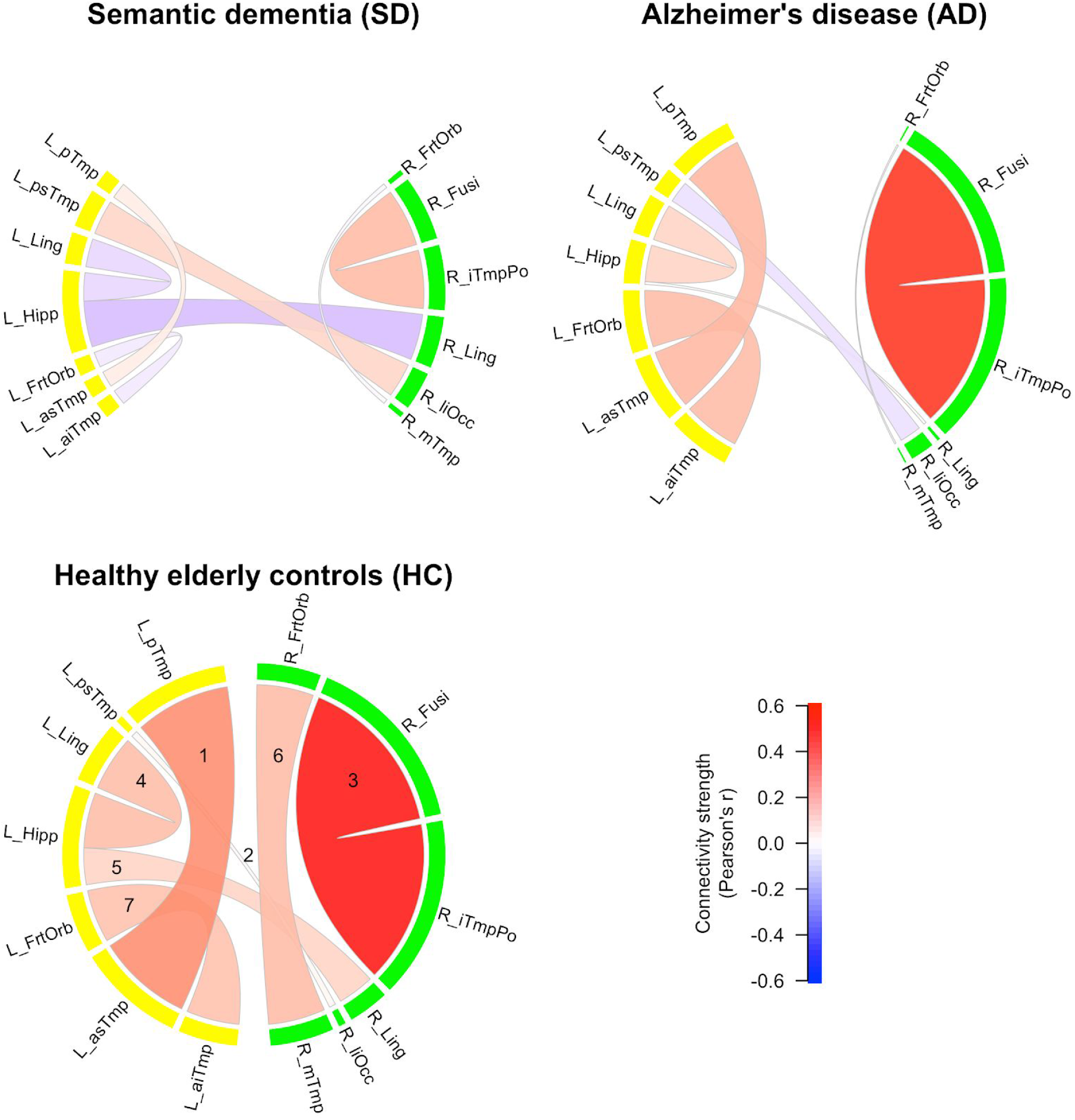
Functional connectivity (FC) strengths of the three groups. Color shades and thickness of the links are proportional to FC strengths; shades of red reflect positive, shades of blue negative strengths. Numbers in HC group indicate edge numbers (see Table 2 for a detailed description of the ROIs). ROIs in the left hemisphere are labeled yellow, ROIs in the right hemisphere labeled green.

We performed a sensitivity analysis to demonstrate the robustness of the results. Age, mean gray matter atrophy in the temporal cortex, and study site as covariates produced statistically significant differences in the same seven edges as reported above. For results of the sensitivity analysis with covariates, see Supplementary Tables 1 and 2.

## Discussion

In this study, we compared functional connectivity between semantic dementia (SD), Alzheimer’s disease (AD), and elderly healthy controls (HC) using a functional parcellation of 66 ROIs of the temporal cortex and hippocampus to investigate intra-temporal connections and connections with contralateral temporal regions. The overall picture that emerges is that between the majority of the significant ROIs, SD demonstrated the most striking decrease in FC. In the AD group, most differences compared to the HC group did not reach significance. We believe that the often described disconnections found in AD were not detected in our study due to the mild progression of the disease in our AD group. One reason could be that the remaining gray matter volume in brain regions typically affected by neuronal degeneration was sufficient to maintain an intact FC to remote areas. In other words, the damage found in mild stages might affect the intra-regional processing in local neuronal populations, whereas the inter-regional (i.e., network) FC would be affected during more advanced AD progression [59]. Future studies will require larger sample sizes to demonstrate smaller changes in FC seen even with mild state impairments.

The most intriguing finding of our study for the SD group was the decreased FC between the left posterior hippocampus and left/ right lingual gyri (edges 4 and 5). These disruptions are characteristic for the neurophysiological basis of the SD patients’ typical symptomatology involving an impaired semantic memory. For instance, Sormaz et al. [60] recently showed a correlation of FC between left the hippocampus and the lingual gyrus with topographic memory, and a correlation of semantic memory performance with FC to the intracalcarine cortex, a finding consistent with our results.

Functional connectivity between the left anterior superior/ middle temporal gyrus/ insula and the left posterior middle/ superior temporal gyrus was decreased in SD compared to HC (edge no. 1). It is important to note that this is the single connection that showed an FC difference between SD and HC exclusively (i.e., a finding specific for the SD-HC group single-comparison while neither AD-HC nor AD-SD were significant). These regions are commonly associated with cross-modal integration (as is the hypothesized semantic hub) of auditory and language processing, as well as the processing of the emotionally relevant context. Hence, this finding might reflect the severe semantic deficits in SD (see table 1) that are manifested by the loss of conceptual knowledge [7].

The single connection that showed increased FC in SD compared to the other groups (edge no. 2) was between the left posterior superior temporal gyrus/ parietal opercular cortex/ planum temporale and the right lateral inferior/ superior occipital cortex. The temporal brain areas that constitute this connection are important for early context integration of acoustically presented words [61], lexico-semantic retrieval [62], and are part of a supramodal semantic network [63]. The occipital ROI of this connection subserves visual integration. Thus, an increased FC between these regions might reflect a functional reorganization that is characterized by supporting language comprehension using more sensory inputs. Moreover, this result indicated a reduced hemispheric functional specialization and perhaps an attempt to pool resources that are spared by the pathological developments in SD.

In comparison with AD and HC, SD patients showed a lower FC between the functional ROI encompassing the right fusiform/ parahippocampal gyri and the right inferior temporal pole. Disruption of this connection (no. 3) can be viewed as SD-typical, as the functional profile of the involved regions conforms to SD symptomatology. In particular, the right temporal pole is crucial for non-verbal (e.g., visual) object recognition, which is a hallmark impairment in SD associated with the loss of semantic knowledge [64,65]. The right fusiform gyrus on the other hand is associated with working memory for faces, face perception, and non-verbal associative semantic knowledge [66–68], and the right parahippocampal gyrus is associated with working memory for object location as well as a function as an episodic buffer [69,70]. In line with this, the patients with SD in the present study showed severe object recognition deficits assessed with BNT and oral picture-naming (table 1), even though no behavioral data about face perception or non-verbal semantic knowledge was available.

Similar to connection no. 3, FC was reduced in SD compared with AD and HC between the ROI comprised of left lingual/ intracalcarine/ precuneus cortex and the ROI including left posterior hippocampus/ thalamus (connection no. 4). This finding is in line with Seeley et al. [14], who reported the medial temporal lobe as part of an SD-vulnerable network. Thus, in addition we showed a possible contribution of the primary visual (intracalcarine cortex), visual memory (lingual gyrus) and self-awareness (precuneus, i.e., default mode network) regions to that semantic network. It might appear surprising that the FC of the AD group was not significantly reduced in this connection, despite the commonly known medial temporal lobe atrophy and the pivotal role of the hippocampus in episodic memory encoding [36,71]. However, functional and anatomical changes do not necessarily overlap, and for instance, stable FC of the left hippocampus in early AD (except with right lateral prefrontal cortex) has been reported previously [29].

Lower FC in SD than in AD (and HC) was also found between the right lingual/ intracalcarine cortex and the left posterior hippocampus/ thalamus (connection no. 5). Therefore, connections between bilateral lingual gyri and the left hippocampus were detected in our HC sample (for illustration, see figure 3, connections no. 4 and 5), whereas either of them were damaged in SD, but not in AD. This supports the recent indication of a hippocampal contribution to the semantic memory network [36]. Because episodic memory is relatively spared in SD, the connections between the left posterior hippocampus and the bilateral lingual gyri might contribute to the semantic memory network. On the other hand, we did not find an expected decrease of FC in connection no. 4 in AD, although the precuneus and hippocampus contribute to episodic memory, which is typically impaired in AD. However, we have to bear in mind that our analysis was restricted to temporal lobe FC and thus did not cover the entire episodic memory network, including brain regions located in frontal and parietal lobes. In addition, no episodic memory data were available for the entire sample of our study. Future studies should investigate additional ROIs from the aforementioned areas using larger sample sizes to tackle the increased number of connections and multiple testing correction that are associated with larger networks.

The only FC reduction common to both SD and AD compared with HC was found in connection no. 6. The functional role of the involved regions suggests an association with a frequently observed clinical presentation of AD and SD characterized by apathy and agitation, associated with the right orbitofrontal cortex [72,73], and impairments in social behavior related to the right anterior temporal lobe [74]. According to Olson et al. [75], social knowledge is part of semantic memory and involves memory about people including biographical information. Nonetheless, caution is advised with comparing social or semantic deficits between AD and SD; both symptoms have different onsets or severities within disease stages, as well as different characteristics. Furthermore, we did not have data on social behavior or apathy/agitation of our patients. Regardless, we added a common pathway to the crossroad described by La Joie et al. [36]. They suggested that the hippocampus is a converging hub of an (AD-affected) episodic and a (SD-affected) semantic network. Accordingly, our data indicated that besides a shared damaged hub in AD and SD, the functional connection between the right anterior middle/ superior temporal gyri and the right orbitofrontal cortex might be a second candidate for the neuropathology shared in both clinical populations.

The final significant connection (no. 7) of the present study was found between the left orbitofrontal cortex and the left anterior inferior and middle temporal gyri. The literature suggests a functional role of this connection in deficient socioemotional abilities that are found predominantly in the behavioral variant of FTD [76], and in higher level object representation, involving language and auditory processing. Unlike in connection no. 6, the AD group did not show an impaired FC of the orbitofrontal regions with the ipsilateral temporal cortex. Thus, one might speculate about a bilateral breakdown of orbitofrontal to temporal connections in SD, which might be related to the severity of the semantic deficit.

We also want to address some limitations of our study. Even though the overall sample size is large, the sample sizes of the three subgroups are considered small (16–23 individuals). Larger studies need to be conducted, however, this is especially challenging for SD given its low prevalence. Therefore, we pooled two SD samples from two different sites with different scanners; however, most individuals of the SD group were from the Shanghai site which is a design problem and potential confound. Moreover, the diagnostic criteria of the two sites for SD were not identical. The data from different MRI sites may have different noise levels such as thermal noise, physiological noise, and motion [52,77]. These artifacts are often global in nature, and global signal regression (GSR) can successfully remove these and standardize the data between sites and across individuals. That means the distribution of correlation coefficients is shifted from predominantly positive to mean zero with positive and negative coefficients. The negative correlations can appear as significant anti-correlations and need to be interpreted with caution, however, GSR can also improve the specificity of positive correlations [78]. Importantly, in this study, we do not interpret absolute negative correlations and solely compare *relative differences* in correlations between groups. Moreover, interpretation of group differences between AD and SD should take into account that the two dementia groups were not matched for disease stage (i.e., SD showed more severe deficits than AD). Likewise, more symptom specific behavioral scores (other than MMSE) could have aided an in-depth interpretation of altered FC edges in the patient groups. Lastly, our analysis did not cover all brain regions potentially relevant for AD and SD. However, the choice to limit the scope to the temporal lobe has three reasons: first, the distinct temporal lobe atrophy is crucial for AD and SD differentiation. Second, the temporal lobe is pivotal in both semantic and episodic memory functions. Third, the definition of ROIs within the temporal lobe, even though using an arbitrary selection threshold of 5% (or more) of overlap of the ROIs with any temporal structure, may be altogether less arbitrary and biased compared to subjectively selecting ROIs based on expectations and literature.

To summarize the main findings of our study, the cohort of patients with SD yielded a number of distinct ipsilateral and contralateral connections of the temporal lobe that showed a significant reduction in FC. These connections included the regions on which our predictions were based on (i.e., hippocampus, fusiform gyrus, and temporal pole). Two functional connections were intriguing due to their distinctiveness from the other groups: the first was the connectivity breakdown between left posterior hippocampus and bilateral lingual gyri, likely reflecting the neuronal underpinning of semantic memory loss. Second, a bilateral disruption of connectivity between temporal and frontal lobes was found. This aligns well with the pathophysiology within the FTD spectrum and especially with SD.

FC in AD was relatively intact compared to SD, which contradicted our hypothesis. The only connection with significantly reduced FC encompassed the right orbitofrontal cortex and the right anterior temporal lobe (no. 6), which we identified as an AD/SD-common pathway. Additionally, figure 2 illustrates that our AD group had a lower FC than the HC in connections no. 1, 2, and to a smaller extent no. 5 which all missed significance. These FC signatures in the AD group could be attributed to their mild stage of symptom progression (MMSE of 24.5), and potentially an early marker of the disease, but larger and longitudinal studies are needed.

Following the “cortically distributed plus semantic hub” theory, several connections were found to be significantly altered in the present study, which affected the anterior temporal lobe – semantic hub – regions (no. 1, 3, 6, and 7). Moreover, their counterparts were partly localized in the modality-specific regions described by Patterson et al. [5], but also in orbitofrontal regions. This agreement of our results with the arguments in Patterson et al. [5] supported the “distributed plus hub” theory, because we found altered FC in connections between the hub and the modality-specific regions. Taken together, this study presents an alternative concept to investigate the understanding of distinct pathophysiological changes in AD and SD that are related to disruptions of functional networks in the temporal lobe. The unique aspect of our study was the definition of ROIs based on functional brain segregation rather than anatomy for FC analysis. Due to the comparably strict statistical approach and the predefined choice of ROIs, our study provided a fine-grained overview of FC aberration related to temporal lobe function in AD and SD. However, comparability was limited owing to different study sites using partially different diagnostic criteria and data acquisition procedures, thus, our findings ideally motivate future studies for replication with identical MRI acquisition parameters and same MRI scanner types.

## Data Availability

Raw imaging data can be requested from the corresponding author. Aggregated data and analysis scripts to generate all results and figures are available at OSF (https://osf.io/t4jnv/).

## Acknowledgments

This work was supported by the Swedish Alzheimerfonden, the Swiss Synapsis Foundation, and the University of Bern, the Beijing Natural Science Foundation (7182088), and the NSFC (31872785). Simon Schwab acknowledges funding from the Swiss National Science Foundation (SNSF, No. 162066 and 171598).

## Abbreviations

AD: Alzheimer’s disease
FC: functional connectivity
HC: healthy controls
ROI: region of interest
SD: semantic dementia

---

**Supplementary Figure 1.**
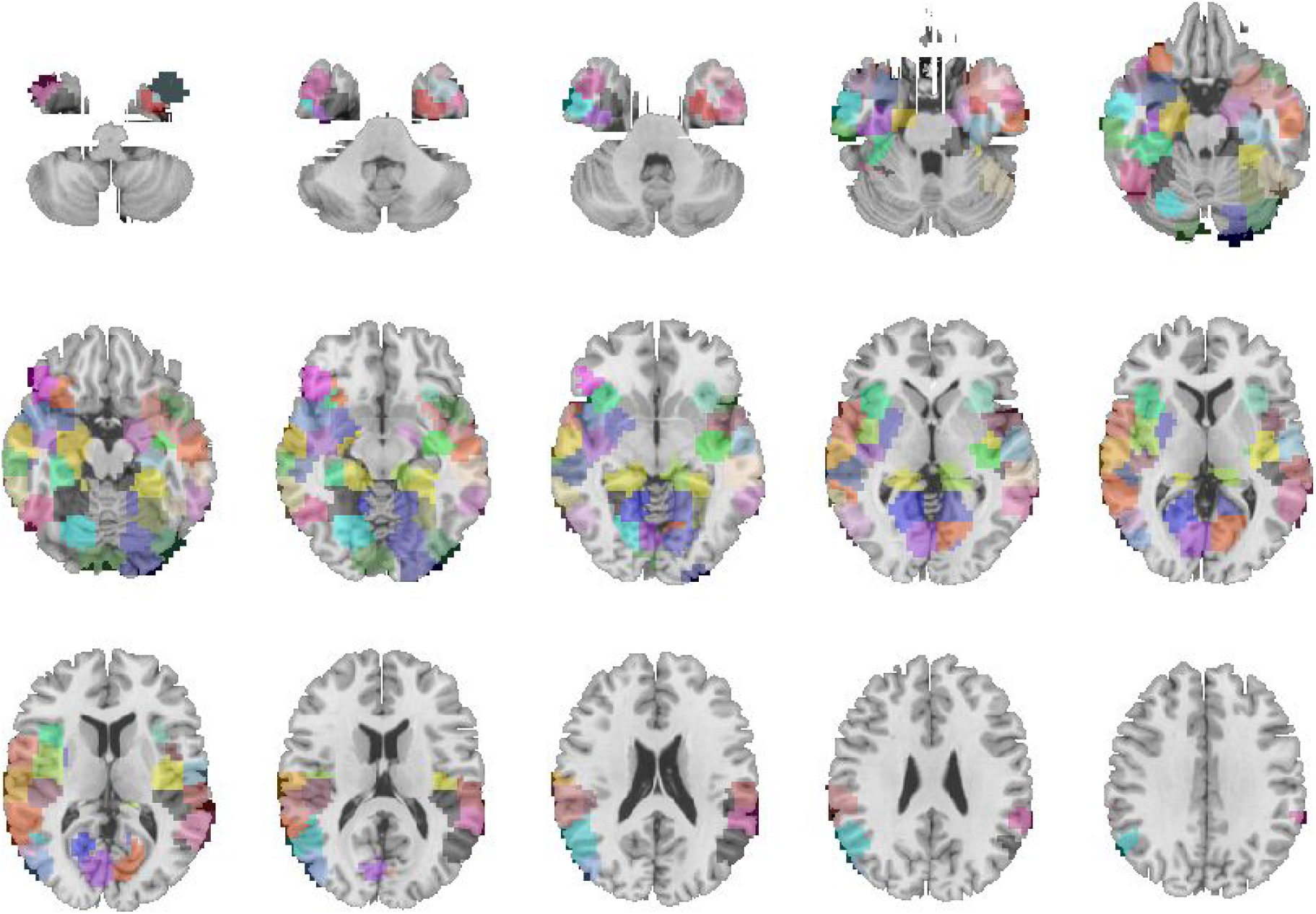
Sixty-six ROIs from a functional parcellation by Craddock [38] (k = 200) that were selected based on their overlap with structures in the temporal cortex, the hippocampus, and the amygdala.

**Supplementary Figure 2.**
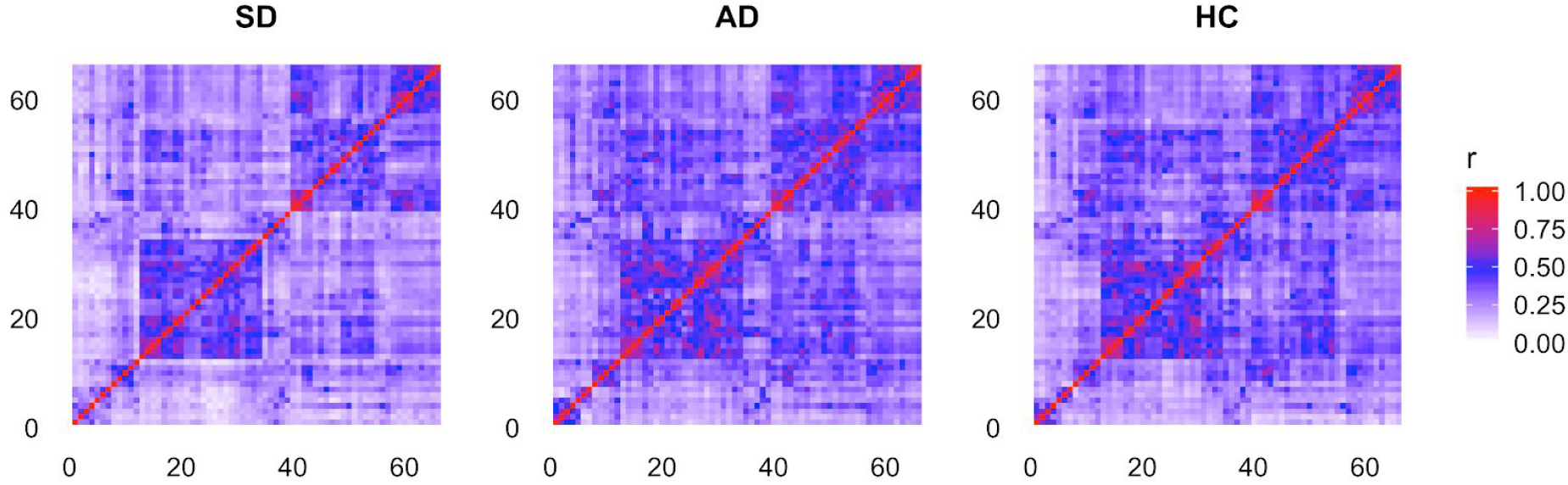
Mean connectivity strengths shown as correlation matrix (Pearson’s r) for the 66 ROIs from the temporal cortex and the three groups (SD, semantic dementia; AD, Alzheimer’s disease; HC, healthy elderly controls).

**Supplementary Figure 3.**
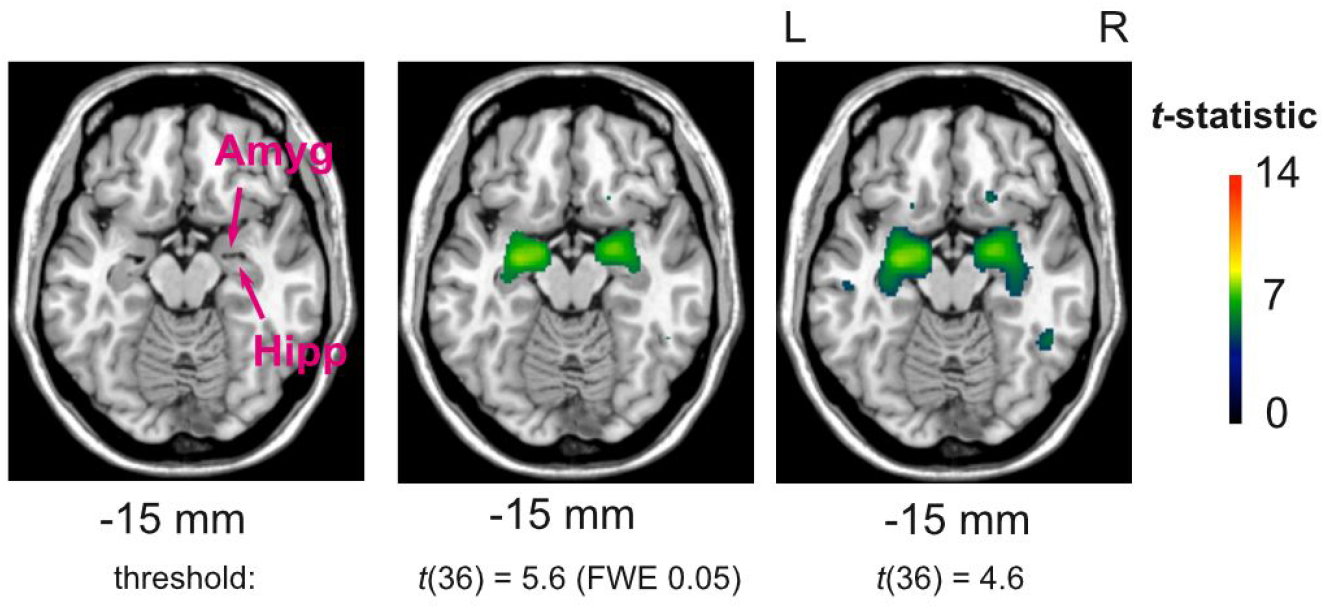
Using a more liberal threshold to increase power reveals reduced gray matter atrophy not only in the amygdala but also in the hippocampus in the Alzheimer’s disease patients.

**Supplementary Table 1.** Sensitivity analysis using three additional covariates, age, scanner site and gray matter atrophy in the temporal cortex. The same seven functional connections that demonstrated significant group differences.

| Edge no. | ROI no. | ROI no. | Region | Region | F-value | FDR adj. p-value |
| --- | --- | --- | --- | --- | --- | --- |
| 1 | 129 | 11 | Left anterior superior temporal gyrus / middle temporal gyrus / insular cortex | Left posterior middle temporal gyrus / superior temporal gyrus | 10.72 | 0.041 |
| 2 | 85 | 24 | Right lateral inferior occipital cortex / lateral superior occipital cortex | Left posterior superior temporal gyrus / central opercular cortex / parietal opercular cortex / planum temporale | 12.27 | 0.025 |
| 3 | 198 | 32 | Right fusiform cortex / parahippocampal gyrus | Right inferior temporal pole | 12.20 | 0.025 |
| 4 | 70 | 37 | Left lingual gyrus / intracalcarine cortex / precuneus cortex | Left posterior hippocampus / thalamus | 12.97 | 0.025 |
| 5 | 89 | 37 | Right lingual gyrus / intracalcarine cortex | Left posterior hippocampus / thalamus | 11.28 | 0.032 |
| 6 | 153 | 71 | Right anterior middle temporal gyrus / superior temporal gyrus | Right orbitofrontal cortex | 11.91 | 0.025 |
| 7 | 112 | 72 | Left orbitofrontal cortex / insular cortex | Left anterior inferior temporal gyrus / middle temporal gyrus | 12.47 | 0.025 |

**Supplementary Table 2.** Post-hoc Tukey HSD *p*-values for the three single comparisons (rows) and for each of the seven ROI-pairs that had a significant group effect (columns). Significant values reflect that the group effect was driven by a specific group level contrast. This analysis included the covariates age, mean gray matter atrophy in the temporal cortex, and study site.

| Edge No. | 1 | 2 | 3 | 4 | 5 | 6 | 7 |
| --- | --- | --- | --- | --- | --- | --- | --- |
| AD vs HC | 0.010 | 0.071 | 0.98 | 0.54 | 0.21 | 0.0007 | 1.00 |
| SD vs HC | < 0.0001 | 0.037 | 0.0002 | < 0.0001 | < 0.0001 | < 0.0001 | 0.0003 |
| SD vs AD | 0.066 | <0.0001 | 0.0006 | 0.002 | 0.022 | 0.98 | 0.0003 |

